# Learning in reverse: Dopamine errors drive excitatory and inhibitory components of backward conditioning in an outcome-specific manner

**DOI:** 10.1101/2022.01.10.475719

**Authors:** Benjamin M. Seitz, Ivy B. Hoang, Aaron P. Blaisdell, Melissa J. Sharpe

## Abstract

For over two decades, midbrain dopamine was considered synonymous with the prediction error in temporal-difference reinforcement learning. Central to this proposal is the notion that reward-predictive stimuli become endowed with the scalar value of predicted rewards. When these cues are subsequently encountered, their predictive value is compared to the value of the actual reward received allowing for the calculation of prediction errors. Phasic firing of dopamine neurons was proposed to reflect this computation, facilitating the backpropagation of value from the predicted reward to the reward-predictive stimulus, thus reducing future prediction errors. There are two critical assumptions of this proposal: 1) that dopamine errors can only facilitate learning about scalar value and not more complex features of predicted rewards, and 2) that the dopamine signal can only be involved in anticipatory learning in which cues or actions precede rewards. Recent work has challenged the first assumption, demonstrating that phasic dopamine signals across species are involved in learning about more complex features of the predicted outcomes, in a manner that transcends this value computation. Here, we tested the validity of the second assumption. Specifically, we examined whether phasic midbrain dopamine activity would be necessary for backward conditioning—when a neutral cue reliably *follows* a rewarding outcome. Using a specific Pavlovian-to-Instrumental Transfer (PIT) procedure, we show rats learn both excitatory and inhibitory components of a backward association, and that this association entails knowledge of the specific identity of the reward and cue. We demonstrate that brief optogenetic inhibition of VTA_DA_ neurons timed to the transition between the reward and cue, reduces both of these components of backward conditioning. These findings suggest VTA_DA_ neurons are capable of facilitating associations between contiguously occurring events, regardless of the content of those events. We conclude that these data are in line with suggestions that the VTA_DA_ error acts as a universal teaching signal. This may provide insight into why dopamine function has been implicated in a myriad of psychological disorders that are characterized by very distinct reinforcement-learning deficits.

**Graphical Abstract:** 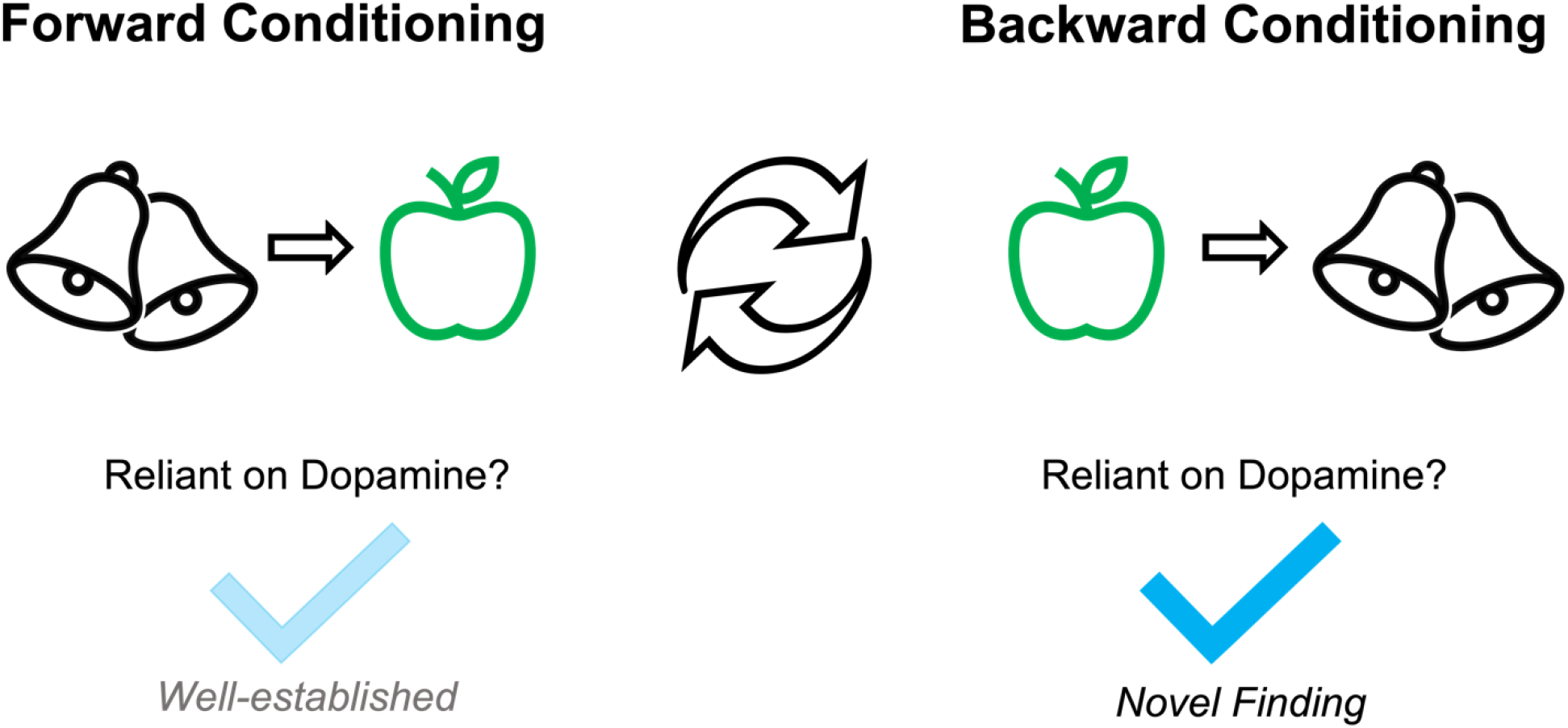

## Results

### Inhibition of VTA_DA_ transients during backward conditioning prevents backward cues from exerting control over instrumental behavior

Early on, studies of associative learning were primarily concerned with understanding the basic mechanisms by which two events—broadly defined—become linked in the brain ^1,2^. It is only recently that a shift has occurred such that major emphasis has been placed on the very specific temporal scenario in which a cue precedes a motivationally-significant outcome (e.g., reward or pain) ^3–7^. Focusing on anticipatory cue→reward learning is advantageous in terms of computational modelling ^5,8–10^ but it leaves many learning phenomena that do not involve this specific temporal order unexplained ^11^.

An example of this trend relates to discovery of the dopamine prediction error. Shortly after it was revealed that dopamine neurons in the midbrain exhibit phasic signals to unexpected rewards^12^, this error signal was interpreted as being governed by computational rules that calculate scalar values in the context of anticipatory cue-reward learning ^12–15^. Consequently, the study of the dopamine prediction error was almost exclusively focused on procedures involving anticipatory cue-reward associations that manipulate scalar value ^16–24^. Only recently have we begun to explore the role of dopamine neurons in more complex paradigms outside of simple cue→reward learning. This work has uncovered that the prediction-error signal is capable of driving anticipatory learning of sensory events that transcend scalar value inherent in rewards, such as an association between two neutral cues ^25–33^. Such findings question the assumption that dopamine neurons are “specialized” for anticipatory reward learning specifically, and whether anticipatory reward learning is “special” more generally.

Backward conditioning—when a reward is *followed* by a cue (reward→cue)—breaks this temporal mold and provides a serious challenge to current computational hypotheses of dopamine function. Backward conditioning can result in both excitatory and inhibitory associations ^34–38^. That is, a backward cue is capable of exciting or inhibiting representation of associated rewards, which motivates the animal towards or away from that specific reward. Here, we tested the necessity of dopamine transients in backward conditioning using an established procedure that combines backward conditioning with Pavlovian-to-Instrumental Transfer (PIT) ^39–41^, which probes for both the specific excitatory and inhibitory components of the association (see Figure S1). This allows us to test whether dopamine neurons are exclusively involved in anticipatory learning, or whether they function as a teaching signal to drive the formation of associations in a broader sense, regardless of whether those associations are anticipatory, inhibitory, or excitatory, and in a manner that transcends scalar value.

Rats expressing Cre-recombinase under the control of the tyrosine hydroxylase (TH) promoter ^42^ received bilateral injections of either inhibitory halorhodopsin (NpHR, AAV5-Ef1a-DIO eNpHR3.0-eYFP, *n* = 9) or control virus that lacks the inhibitory opsin (eYFP, AAV5-Ef1a-DIO-eYFP, *n* = 9) in VTA (see Figure 1). Optic fibers were also implanted bilaterally over VTA. After recovery, rats were food restricted and then received backward training, where two distinct rewards (pellets and maltodextrin solution) were each followed by one of two auditory cues [white noise and clicker (counterbalanced); 8 days, 24 presentations per day]. The pairing of the reward and cue were arranged such that the cue would be presented 10s after the rat entered the magazine to consume the reward. This ensured the cue would be delivered shortly after the rats had consumed the reward. We delivered green light (532nm, 16–18 mW output) into the VTA 500ms before the onset of the cue and continuing for 2s, as we have done previously ^31,43^. We used these parameters to prevent phasic firing at the onset of the backward cue, which would suppress a potential prediction error to the backward cue, without producing a negative prediction error ^27^.

**Figure 1.**
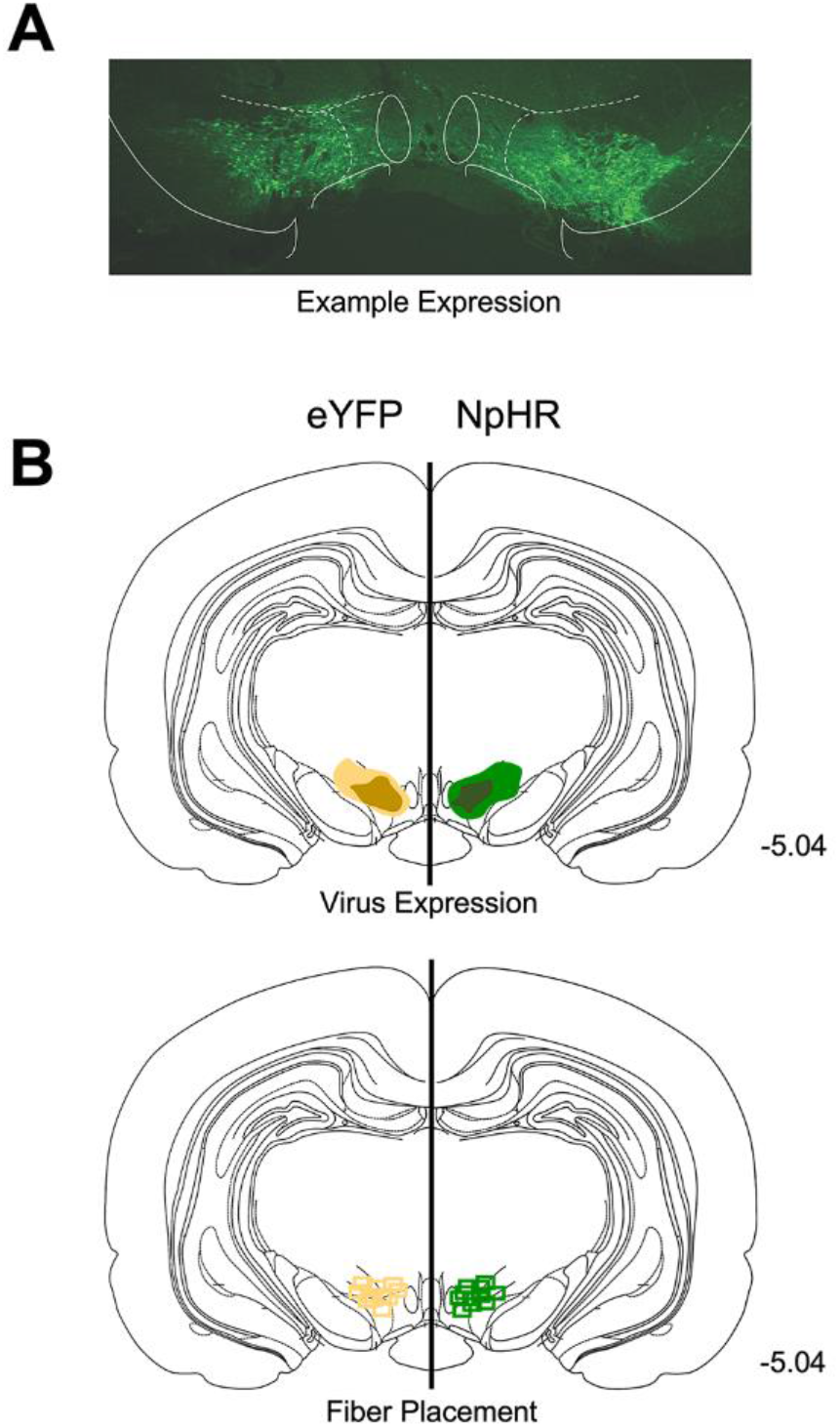
Histological representation of virus expression and fiber placement in TH-Cre rats. A) Neurons in VTA expressing eYFP. B) Unilateral representation of the bilateral virus expression (upper) and fiber placements (lower). Fiber implants (green and yellow squares) were localized in the vicinity of NpHR (green) and eYFP (yellow) expression in VTA.

Responding to the cues decreased over the course of conditioning, in line with other backward conditioning reports ^39–41^, and this was similar across groups (Figure 2A; day: F_7, 112_= 4.593, p = 0.005; group: F_1, 16_ = 0.218, p = 0.647; day x group: F_7, 112_= 0.445, p = 0.741; Figure 2A). Rats then learned to press different levers for the distinct rewards (e.g., left lever→pellets; right lever→solution, counterbalanced), on an increasingly lean random-ratio schedule (CRF, RR5, RR10). All rats acquired the lever-pressing responses with no between-group differences (Figure 2B; day: F_7, 112_ = 650.415, p < 0.001; group: F_1, 16_ = 0.016, p = 0.901; day x group: F_7, 112_ = 1.521, p = 0.227; Figure 2B).

**Figure 2.**
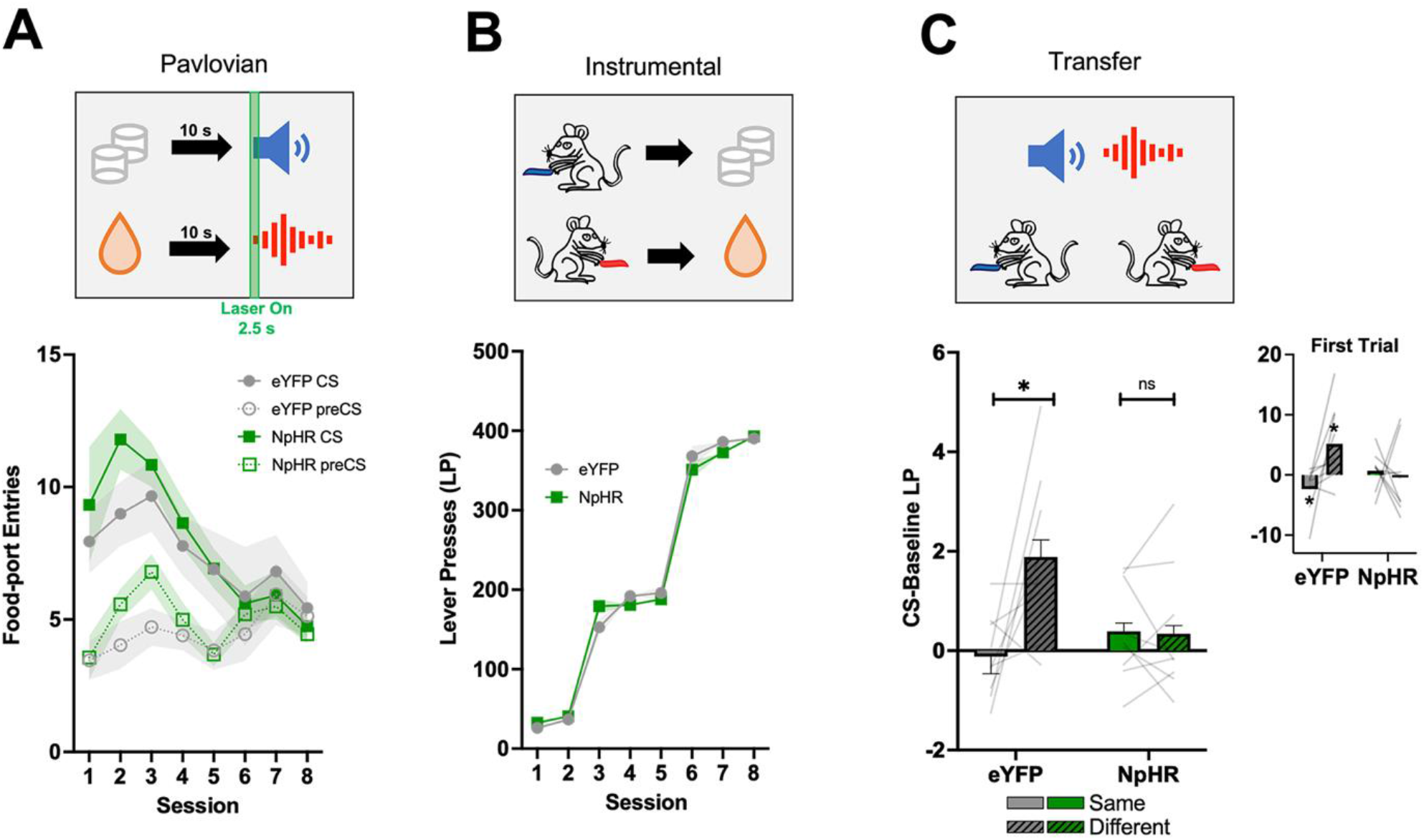
Inhibition of VTA_DA_ transients during backward conditioning prevents backward cues from exerting excitatory and inhibitory control over instrumental behavior. Rates of responding are represented as the number of entries into the food port or lever presses during cue presentation (±SEM), with lines indicating individual data points. **A)** Rats first learned backward relationships between two distinct rewards and two auditory cues (Conditioned stimuli: CSs). The backward cue was presented 10s after the rats entered the magazine to consume the rewards. Here, green light was delivered into VTA at the onset of the backward cue for 2.5s to suppress phasic firing of dopamine neurons without producing a negative prediction error ^27^. Responding during cues decreased over the course of conditioning with no difference between groups. **B)** Rats then learned to make a left lever press to obtain one reward, and a right lever press to obtain the other. All rats acquired the instrumental responses for the rewards, with no difference between groups. **C)** Finally, during the unrewarded PIT test, both levers were made available and the cues were individually presented without rewards (right). During the PIT test, the backward cues biased our eYFP group’s responding away from the associated reward, and towards the lever associated with the different reward. However, our NpHR group showed no change in responding from baseline during cue presentation or bias between the levers. * Indicates significance at p < 0.05.

Finally, rats received a probe test in which both levers were available with no rewards delivered, and the backward cues were presented individually (i.e., the PIT test). The PIT test allows us to examine the nature of the associations that have developed during Pavlovian training. In our eYFP group, backward cues biased lever-pressing away from the associated reward, and towards the alternate reward (Figure 2C; lever x group: F_1, 16_ = 7.054, p = 0.017; simple main effect of lever: F_1, 16_ = 8.318, p = 0.020; see Figure S2 for baseline responding and food-port entries). That is, the pellet-associated backward cue led to rats pressing more for solution, and the solution-associated backward cue led rats to press more for the pellet, in line with previous studies ^39–41^. This shows that the backward cues excite one behavior (lever press for different reward), while also inhibiting the other (lever press for same reward), in a sensory-specific manner. Indeed, on the first trial, responding in our eYFP group to the different lever was significantly elevated from baseline (t_8_ = 2.474, p = 0.038) whereas analyses suggested responding on the same lever was lower than baseline (t_8_ = 5.500, p = 0.050). However, rats in our NpHR group showed no bias on lever responding and were not *elevated or decreased* from baseline lever-press responses (simple main effect of lever: F_1, 16_ = 0.021, p = 0.889; different lever versus baseline on first trial: t_8_ = 0.202, p = 0.845; same lever versus baseline on first trial: t_8_ = 0.669, p = 0.504). Finally, baseline lever press responding did not statistically differ between the two groups, t_16_ = 0.946, p = 0.358 (Figure S2A). Similarly, head entries into the food-port did not differ between groups, t_16_ = 0.480, p = 0.638 (Figure S2B). These findings suggest that inhibition of VTA_DA_ neurons at cue onset prevent the backward cues from exerting any effect over instrumental responding for the paired rewards, in an inhibitory or excitatory manner.

### Inhibition of VTA_DA_ Neurons Prevents Acquisition of the Specific and General Inhibitory Components of Backward Conditioning

There are multiple interpretations that could be made from the failure of our NpHR group to use the backward cues to modulate instrumental performance. We suggest that VTA_DA_ inhibition prevented learning about the excitatory and inhibitory relationships between the rewards and backward cues. However, it is also possible that the NpHR rats still learned the inhibitory associations, but that the cues lacked some aspect of motivational significance that would allow them to exert control over an instrumental response. A second interpretation of the PIT data is that the NpHR rats may have learned the backwards cues were generally inhibitory of rewards. Thus, the performance of the NpHR rats during the PIT test could be interpreted as blanket inhibition of both lever-press responses during the PIT test—though this is unlikely as these rats did not reduce lever-pressing from baseline in the PIT test (see Figure 2C).

To dissociate these accounts, we next taught the same rats two new forward associations with visual cues (e.g., house light→pellets; flashing light→maltodextrin solution; Figure S2). Training these new associations allowed us to investigate the impact of the backward cues on Pavlovian responding when presented in compound with the visual cues in an un-rewarded test session (i.e., a summation test). That is, when presented by themselves the visual cues should elicit high levels of responding because they signal the occurrence of a rewarding outcome. However, when each visual cue is presented in compound with the backward cue that signals the absence of the same outcome (i.e., a congruent compound), responding should be considerably reduced if the auditory cues are inhibitory. As predicted, responding in group eYFP was high when the visual cue was presented individually, while pairing it with the congruent backward cue significantly attenuated responding (Figure 3: Summation test; cue type x group: F_1, 9_ = 11.893, p = 0.007; simple main effect of cue type: F_1, 9_ = 16.975, p = 0.009). However, in the NpHR group, the presence of the backward cue had no impact on responding to the visual cue (simple main effect of cue type: F_1, 9_ = 0.375, p = 0.573). This confirmed that the backward cues possessed inhibitory properties that could influence Pavlovian responding, and that inhibition of VTA_DA_ neurons prevented backward cues from acquiring inhibitory properties.

**Figure 3.**
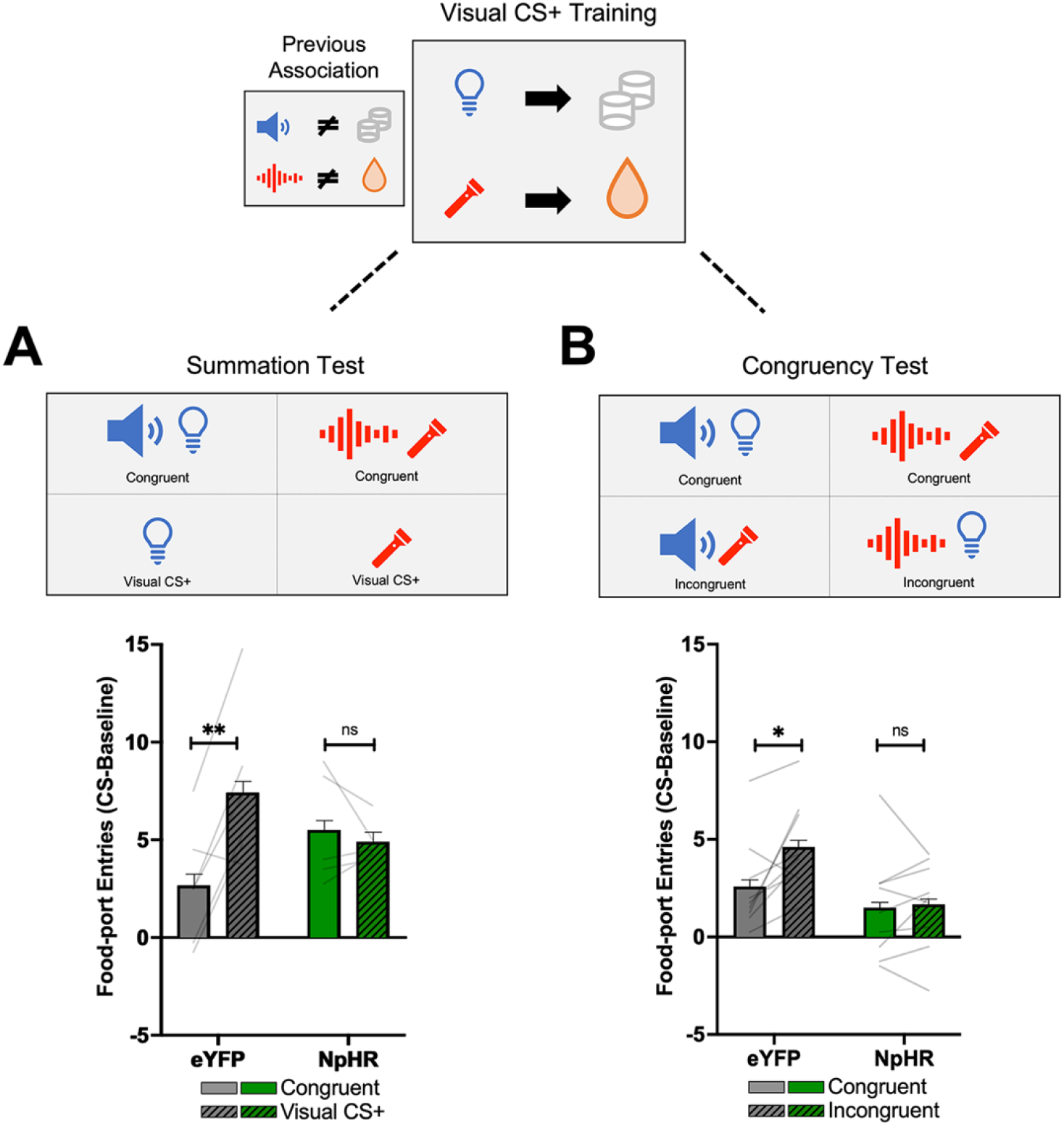
Inhibition of VTA_DA_ transients prevents backward cues from generally and specifically inhibiting Pavlovian responses. Responding is represented as number of entries into the food port during cue presentation (±SEM), with lines indicating individual data points. **Top)** *visual forward training*: To assess the nature of the deficit in the instrumental PIT test, we trained rats with two new forward cue-reward associations with visual stimuli (Figure S3). This allowed us to perform a number of tests with novel audiovisual compounds to investigate the source of the deficit in our NpHR group. **A)** *Summation test:* we tested responding to the visual cue by itself, relative to when it was presented in compound with the backward cue associated with the same outcome (i.e., congruent compound). If the backward cue is inhibitory, responding should be reduced on congruent trials relative to trials with the visual cue alone. Indeed, this is what we observed in the eYFP group. In contrast, the NpHR group showed the same high levels of responding to the visual cue whether or not it was presented in compound with the backward cue. **B)** *Congruency test:* The previous test indicates the backward cues are inhibitory when paired with the same outcome, but did not test whether those cues possess specific or general inhibitory properties. To test this, we presented the visual cues in compound with the auditory cue predicting the same (congruent) or different (incongruent) reward. In the eYFP group, rats responded less on congruent relative to incongruent trials, suggesting the backward cues were specifically inhibitory. Again, there was no effect of the backwards cues on responding to the visual cues in the NpHR group. *Indicates significance at p < 0.05, **Indicates significance at p < 0.01.

While the summation test above shows that VTA_DA_ inhibition prevents animals from learning the inhibitory component of backward cues in a Pavlovian procedure, they cannot speak to whether the backward cues generally or specifically inhibit Pavlovian responding in either the NpHR or eYFP rats. This is because we only presented a compound where both cues were associated with the same outcome and thus do not know if a backward cue presented in compound with a visual cue associated with the different outcome would similarly inhibit responding in a general fashion. A congruency test was used to tease apart the general versus specific nature of the inhibitory relationship that our NpHR group failed to learn. Specifically, just as we had previously presented in compound backward and forward cues associated with the same outcome (i.e., congruent compound), we could also present in compound backward and forward cues associated with different outcomes (i.e., incongruent). If the inhibitory relationship is specific, congruent compounds should show reduced responding relative to incongruent compounds. However, if the inhibitory relationship is general, there should be no difference between congruent and incongruent compounds. In our eYFP group, we observed a reduction in responding on congruent relative to incongruent compound trials (Figure 3: Congruency Test; compound x group: F_1, 16_ = 4.571, p = 0.048; simple main effect of compound: F_1,16_ = 8.790, p = 0.018). In contrast, rats in group NpHR showed no difference in Pavlovian responding during congruent versus incongruent trials (simple main effect of compound: F_1,16_ = 0.096, p = 0.765), confirming they had not learned the specific inhibitory associations with the backwards cue, and it was not a more general deficit in using the Pavlovian cues to exert control over instrumental behavior.

### Inhibition of VTA_DA_ Neurons at Cue Onset in Forward Conditioning Does Not Prevent Learning or Make Cues Aversive

Our prior results showed that brief optogenetic inhibition of VTA_DA_ neurons at cue onset in backward conditioning prevented rats from learning the excitatory and inhibitory components in backward conditioning, which we would interpret as indicating the dopamine prediction error is a broad teaching signal that transcends both scalar value and anticipatory associative structures. However, it is possible that inhibiting VTA_DA_ neurons at cue onset somehow made these cues aversive, or simply reduced their salience so that they could not be learned about. To test this, we taught all rats new forward relationships between two novel auditory cues (siren and tone) and two distinct food rewards in a novel context. We delivered green light (532 nm, 16–18 mW output) to VTA_DA_ neurons at cue onset for one of the auditory cues but not the other (counterbalanced), using the same inhibition parameters as backward conditioning (i.e. 2.5s inhibition at cue onset). We observed no difference in acquisition between the cue with laser on versus the cue with the laser off in either group (Figure 4A; day: F_7, 112_ = 2.741, p = 0.060; laser: F_1, 16_ = 0.947, p = 0.345; group: F_1, 16_ = 0.079, p = 0.782; day x group: F_7, 112_ = 0.246, p = 0.845; day x laser: F_7, 112_ = 1.266, p = 0.291; laser x group: F_1, 16_ = 2.051, p = 0.171; day x laser x group: F_7, 112_ = 0.522, p = 0.734). However, in the NpHR group, the cue with the laser on showed a small, but statistically non-significant, retardation of acquisition, approximately replicating the results of Morrens et al ^44^ (simple main effect of laser status: F_1,16_ = 3.940, p = 0.082; Figure 4A). Despite this, responding during the two cues was virtually indistinguishable after the initial sessions, and an extinction test after the completion of training revealed no between-group or within-group differences in responding (Figure 4B; laser status: F_1, 16_ = 0.236, p = 0.634; group: F_1, 16_ = 0.011, p = 0.916, laser status x group: F_1, 16_ = 0.006, p = 0.937). These results suggest that VTA_DA_ inhibition at cue onset does not prevent learning about the cue-reward association. Thus, the results from the previous studies cannot be explained by VTA_DA_ neuronal inhibition reducing the salience of the cues to the extent that they cannot be learned about or making them in some way aversive.

**Figure 3.**
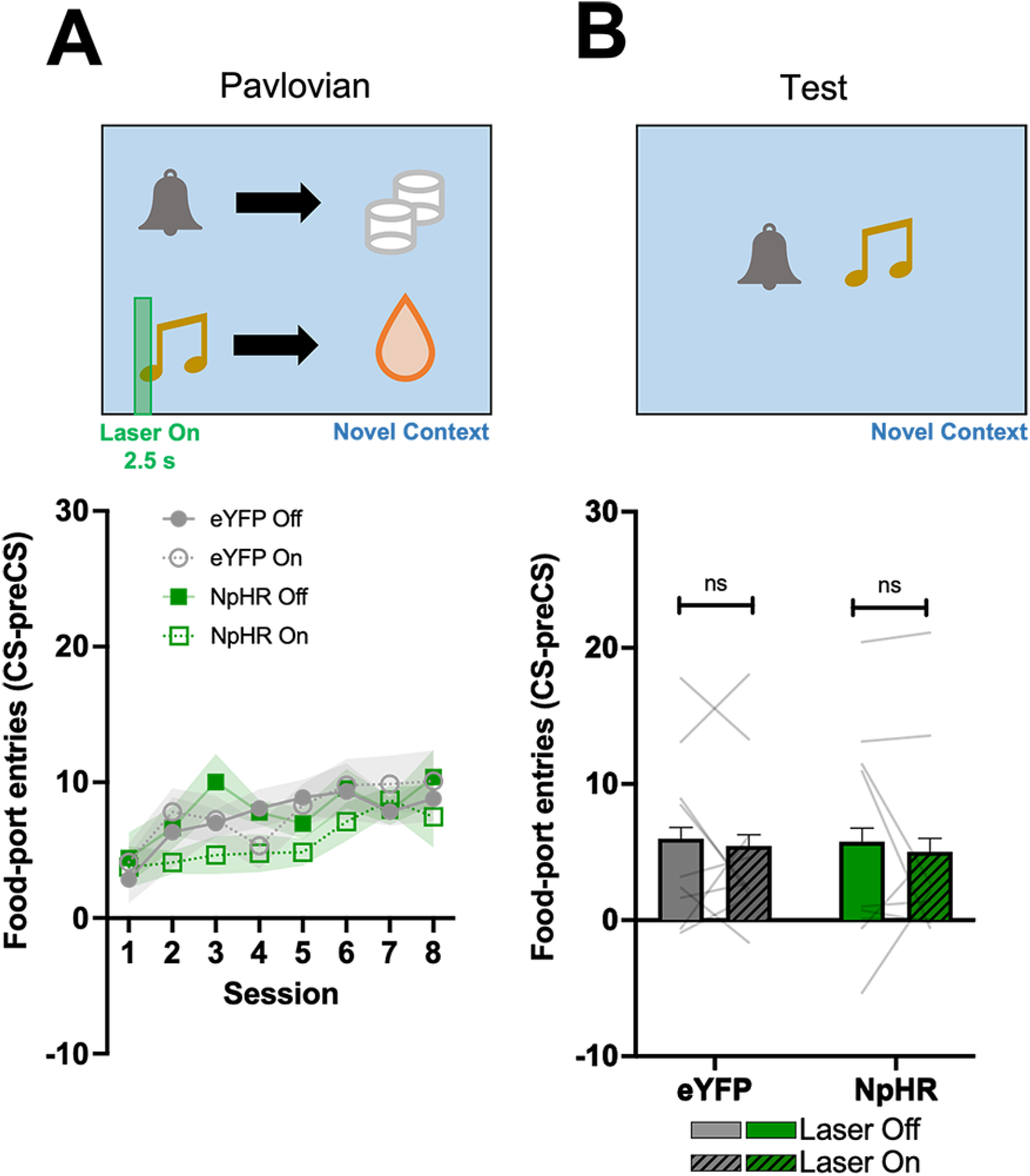
Inhibition of VTA_DA_ transients at cue onset in forward conditioning does not impair learning. Responding is represented as number of entries into the food port during cue presentation (±SEM). To ensure our findings could not be the result of VTA_DA_ inhibition at cue onset causing the backward cues to become aversive or reducing their salience, we taught rats novel auditory cue-reward associations with VTA_DA_ inhibition at cue onset. **A)** Rats learned forward relationships with two novel auditory cues, one of which received light delivery into VTA at cue onset. Pavlovian training progressed normally for both cues in group eYFP, with a non-significant reduction in responding to the NpHR group at the beginning of training. **B)** We then tested responding to the auditory cues by themselves without laser inhibition. There were no differences in responding between groups, or between cues. These results suggest VTA_DA_ inhibition at cue onset does not prevent learning.

## Discussion

These data show that backward reward→cue associations can modulate instrumental behavior in an excitatory, inhibitory, and outcome specific manner. Further, inhibition of VTA_DA_ neurons at the onset of the backward cue to suppress phasic firing of dopamine neurons prevents learning of these backward associations. We also ruled out the possibility that inhibiting VTA_DA_ neurons at cue onset simply prevents learning by reducing cue salience. These data are consistent with recent work implicating phasic activity in VTA_DA_ neurons in learning outside the context of scalar values ^25–33^, and extend this research in critical ways.

Canonical models ^10,12–15,45^ of the dopamine prediction error has restricted these neurons to anticipatory cue-reward learning, via the backpropagation of scalar value to a reward-predictive cue. However, our data show that VTA_DA_ transients are necessary for the excitatory and inhibitory components of backward conditioning in a manner that entails specific knowledge of the identity of the events. This comes at a time when there is mounting evidence that the dopamine error facilitates far more complex learning than that afforded by the backpropagation of scalar value ^46,47^. For example, VTA_DA_ transients are necessary and sufficient for learning associations between two neutral cues (e.g., tone→light), and VTA_DA_ neurons achieve this without making the neutral cues valuable in and of themselves ^27,28,31,48^. Similarly, artificially inducing dopamine prediction errors during cue-reward learning allows the cue to evoke a detailed representation of the reward ^49^. Results like these and others ^25–30^ suggest VTA_DA_ neurons are capable of producing an error that facilitates “model-based” learning, which refers to an ability to associate (and predict) sensory representations of events. However, even an error signal that facilitates model-based learning cannot fully explain our results with backward conditioning. This is because model-based accounts still ultimately rely on value back propagating to earlier predictors of reward, albeit in the context of more complex associative structures, whether inferred or directly experienced ^8,50^.

How should we interpret the necessity of VTA_DA_ neurons in backward conditioning? The most parsimonious explanation of our data and other recent findings is that VTA_DA_ neurons are computing prediction errors between contiguously-occurring events. Thus, regardless of if the events are two contiguously-occurring cues (as in sensory preconditioning ^31^ and second-order conditioning ^43^) or other sensory events, VTA_DA_ neurons might be sending errors that reflect a mismatch between sensory expectations and events. That is, it could be considered a more general sensory prediction error, that serves to reduce the presence of prediction errors in our everyday sensory experience, which sometimes involves events that possess value (like rewards). Indeed, the original Rescorla-Wagner model ^3^, which serves as the basis for Temporal Difference Reinforcement Learning (TDRL) algorithms, is agnostic towards whether prediction errors are value-based or more cognitive like we are now suggesting. Such a stance would argue that VTA_DA_ neurons are contributing to learning in ways more closely aligned with historical interpretations of associative learning ^2^ and less with modern TDRL-centric interpretations.

The implications of dopamine acting as a more universal teaching signal are profound. First, if dopamine contributes to mentally linking contiguously-occurring events, rather than for predicting rewards (either proximally or distally), it would explain why it has been found to be necessary for higher-order conditioning ^31,43^, and also places dopamine at the center of many complex forms of cognition (e.g., spatial and causal reasoning) ^51^. Ultimately, this may have important implications in pathologies characterized by abnormal dopaminergic functioning (e.g., schizophrenia and addiction). Indeed, an excess of subcortical dopamine (a trademark of schizophrenia) would be expected to be correlated with an excess in learning relationships between potentially irrelevant events—which could result in hallucinogenic or delusional experiences ^52–57^. To expand, not all co-occurring events need be associated, and there are also regions (e.g., lateral hypothalamus) whose function appears to be opposing the learning of relationships that do not immediately predict rewards ^58,59^. Such findings situate the VTA_DA_ prediction error at the center of a dynamic system whose main function is to direct learning in one way or another via distinct circuits, depending on current context or motivational state, and past experience. Future research will tell how far we can push the boundaries of dopamine’s involvement in learning and cognition.

## EXPERIMENTAL MODEL AND SUBJECT DETAILS

### Subjects

18 transgenic Long-Evans rats (8 Female, 10 Male) expressing Cre-recombinase under the control of the tyrosine hydroxylase promoter were used in this study. Rats were randomly allocated to groups and matched for age and sex. Rats were maintained on a 12-h light–dark cycle, where all behavioral experiments took place during the light cycle. Rats had ad libitum access to food and water unless undergoing the behavioral experiment during which they received sufficient chow to maintain them at ~85% of their free-feeding body weight. All experimental procedures were conducted in accordance with the UCLA Institutional Animal Care and Use Committee.

## METHOD DETAILS

### Surgeries

Surgical procedures have been described elsewhere^31^. Briefly, rats received bilateral infusions of 1.0-2.0 μL of AAV5-EF1α-DIO-eYFP (n = 9) or eNpHR3.0-eYFP (n = 9) into the VTA at the following coordinates relative to bregma: AP: −5.3 mm; ML: ± 0.7 mm; DV: −6.5 mm and −7.7 (females) or −7.0 mm and −8.2 mm (males). Virus was obtained from Addgene. During surgery, optic fibers were implanted bilaterally (200-μm diameter, Thorlabs, CA) at the following coordinates relative to bregma: AP: −5.3 mm; ML: ± 2.61 mm and DV: −7.05 mm (female) or −7.55 mm (male) at an angle of 15° pointed toward the midline.

### Apparatus

Behavioral sessions were conducted in identical sound-attenuated conditioning chambers (Med Associates, St. Albans, VT). The chambers contained 2 retractable levers that could be inserted to the left and right of a recessed food delivery port in the front wall when triggered. A photobeam entry detector was positioned at the entry to the food port. The chambers were also equipped with syringe pumps to deliver 15% maltodextrin solution in 0.1 ml increments through a stainless steel tube into a custom-designed well in the food port and a pellet dispenser to deliver a single 45-mg sucrose pellet (Bio-Serv, Frenchtown, NJ). Both a tone and white noise generator were attached to individual speakers on the wall opposite the lever and magazine. A 3-watt, 24-volt house light mounted on the top of the back wall opposite the food cup and two white lights were mounted above the levers and served as visual cues.

### Backward Pavlovian Training

Rats received 8 consecutive days of Pavlovian conditioning. Outcomes (sucrose pellet or maltodextrin solution) were delivered into the food port, and auditory cues (clicker or white noise) were played 10 s following the first entry into the magazine. Outcome-cue relationships were fully counterbalanced. Cue duration varied from 2-58 s with an average of 30 s. Data are presented as average entries per minute. Variable cue duration was chosen to stay consistent with the procedure described elsewhere ^39–41^ and because variable cue length helps promote instrumental responding at test by preventing the animal from timing the delivery of the outcome. Stimuli were presented 12 times each in a pseudorandom order with a variable inter-trial-interval (ITI) ranging from 80-190 s with an average of 125 s. Rats received three reminder sessions of this training; reminder 1 occurred after instrumental conditioning, reminder 2 occurred after PIT test, and reminder 3 occurred after the incongruent/congruent test.

### Instrumental Training

Rats received 8 consecutive days of Instrumental conditioning. Each day consisted of two training sessions separated by at least 3 hours. In each session, left or right lever was extended for 30 minutes or until 20 outcomes had been received. Lever and outcome relationships were fully counterbalanced as was the time of day (early vs late) for each session. Lever pressing was continuously reinforced for the first 2 days of training, reinforced on a random ratio 5 schedule for days 3-5, and reinforced on a random ratio 10 schedule for days 6-8. Rats received a reminder RR10 session in between the two PIT tests. Data are presented as total number of lever presses per session/day.

### Transfer Test

Rats received 2 transfer test sessions. The sessions were separated by 2 rest days and one RR10 instrumental reminder session. The data is collapsed between the two days and a 2 (Day 1 vs Day 2) x 2 (Same-Baseline vs Different-Baseline) x 2 (eYFP vs NpHR) mixed measures ANOVA revealed no significant effect of day: F_1, 16_ = 2.373, p = 0.143, no interaction between day and group: F_1, 16_ = 0.240, p = 0.631, nor interaction between day and lever: F_1, 16_ = 0.565, p = 0.463. At the start of the session, both levers were extended for 8 min to allow for extinction to the levers. All rats then received the following order of stimulus presentation: white-noise, clicker, clicker, white-noise, clicker, white-noise, white-noise, clicker, as is standard in the field ^39–41^. Thus, each cue was presented 4 times for 60 s. Because cues are counterbalanced relative to the rewards they predict, the order of cue presentation is also counterbalanced in the above order. Lever pressing during the cue is subtracted from a 60 s baseline (average of lever pressing made to both levers prior to each cue presentation). This gives us a measure of how much rats increase (or decrease) responding from baseline during the cues. Data are presented as average lever presses-baseline per minute. Trials were separated by a fixed ITI of 180 s.

### Forward Conditioning with Visual Cues

Rats received 3 consecutive days of Pavlovian training where a visual cue (house light or flashing white lights) predicted the occurrence of an outcome (sucrose pellet or maltodextrin solution). Visual cues were randomly presented 15 times each for a fixed duration of 30 s and immediately terminated with the delivery of the outcome. Responding during the visual cue is measured relative to the number of entries made 30 s before the cue was presented (CS-preCS). Data are presented as average entries per minute. Trials were separated by a variable ITI ranging from 130-230 s with an average of 180 s. Rats received two consecutive reminder sessions of this training after completing the congruency test session and before the summation test.

### Congruency Test

Rats received a single test session responding to congruent/incongruent audiovisual compounds presented in extinction. Four unique compounds (2 congruent and 2 incongruent) were presented four times each. Compounds were presented in the following order: clicker_flash, noise_house, noise_flash, clicker_house, noise_house, clicker_flash, clicker_house, noise_flash. Compounds were presented for a total of 30 s and were measured relative to responding made 30 s prior to compound presentation. Data are presented as average entries per minute. Trials were separated by a variable ITI ranging from 130-230 s with an average of 180 s.

### Summation Test

A subset of rats (N=11) received a single summation test in which the visual cues were presented by themselves or in compounds with the specific auditory cue associated with the same outcome (congruent compound). Each visual cue and audiovisual compound was presented 4 times each for a total of 16 trials. Order of presentation was pseudo-randomly counterbalanced. Cues were presented for a total of 30 s and are measured relative to responding made 30 s prior to compound presentation. Data are presented as average entries per minute. Trials were separated by a variable ITI ranging from 130-230 s with an average of 180 s.

### VTA_DA_ neuronal inhibition at cue onset in forward conditioning

Rats received 8 consecutive days of Pavlovian training in a novel context where novel auditory cues (siren and pure tone) predicted the occurrence of an outcome (sucrose pellet or maltodextrin solution). Auditory cues were randomly presented 15 times each for a fixed duration of 30 s and immediately terminated with the delivery of the outcome. Laser light was delivered for 2.5s beginning 0.5s before cue onset for one of the two cues (counterbalanced). Responding during the cues was measured relative to the number of entries made 30 s before the cue was presented. Trials were separated by a variable ITI ranging from 130-230 s with an average of 180 s. After 8 days of conditioning, rats received a single test session in extinction where each stimulus was presented 8 times without laser delivery. Stimulus presentation was pseudo-randomly ordered and fully counterbalanced. Auditory cues were presented for a total of 30 s and are measured relative to responding made 30 s prior to cue presentation. Trials were separated by a variable ITI ranging from 130-230 s with an average of 180 s. Data are presented as average entries per minute.

### Histology

The rats were euthanized with an overdose of carbon dioxide and perfused with phosphate-buffered saline followed by 4% paraformaldehyde (Santa Cruz Biotechnology Inc.). Fixed brains were cut in 20-μm sections, and images of these brain slices were acquired and examined under a fluorescence microscope (Carl Zeiss Microscopy). The viral spread and optical fiber placement (Figure 2A and 2B) were verified and later analyzed and graphed using Adobe Photoshop.

### Data collection and statistics

Data was collected using Med-Associates automated software and the text file output were analyzed using MPC2XL (Med Associates, St. Albans, VT). Repeated Measures Analysis of Variance (ANOVA) were used to assess training and test data in JASP (version 0.15). Simple main effects were used to follow up on significant interactions and assess the effect of lever (Same vs Diff) on each group (eYFP vs NpHR), the effect of compound type (Incongruent vs Congruent) on each group, and the effect of cue type (Visual CS+ vs Compound) on each group. One sample T-tests were used to measure responding relative to baseline (expected value = 0). All data were tested for normality and analyses that did not pass this criterion were adjusted using a Greenhouse-Geisser (Repeated Measures) or Wilcoxon (T-test) correction. For instances in which a Greenhouse-Geisser correction was used, the adjusted p value is reported but degrees of freedom are reported in their uncorrected form. Pilot data (n=11) presented in the supplementary material revealed the effect of lever on the PIT test was very large, *η^2^* = 0.519 or *f* = 1.039 using the formula (*f* = sqr(*η^2^* / (1 - *η^2^*)). A power analysis conducted in G*power (version 3.1) revealed 8 participants would be necessary to discover a similarly sized effect with 90% power (between measurement *r* = 0.074). Thus, we were well powered to detect a main effect of lever in our initial PIT test with 9 participants per group.

### Pilot Study

A pilot study was conducted in wild-type rats (n=11) to confirm successful influence of backward conditioning on PIT and to replicate the procedure described elsewhere ^39–41^. The procedure was identical to that described in the Pavlovian, Instrumental, and Transfer Test sections in those manuscripts (Figure S2A). Responding to both the pellet and maltodextrin cue decreased over the course of conditioning and there was no difference between cues (day: F_7, 70_ = 3.531, p = 0.003; reward: F_1, 10_ = 0.008, p = 0.931; day x reward: F_7, 70_ = 0.821, p = 0.573; Figure S1A). Rats then learned to press different levers for the distinct rewards on an increasingly lean random-ratio schedule (Figure S1B). All rats acquired the lever-pressing responses with no differences between the rewards (day: F_7, 70_ = 1321.052, p < 0.001; reward: F_1, 10_ = 1.051, p = 0.329; day X reward: F_7, 70_ = 0.992, p = 0.444; Figure S1B). Finally at test, both levers were extended and the backward cues were presented sequentially. Backward cues biased lever pressing towards making the opposite lever press relative to baseline (lever: F_1, 10_ = 10.809, p = 0.008, *η^2^* = 0.519; Figure S1C).

### Forward Conditioning with Visual Cues

All rats readily learned forward relationships between visual cues and rewards (described in detail in Methods) with no difference between groups (day: F_4,64_ = 30.989, p < 0.001; group F_1,16_ = 0.466; p = 0.504, day X group: F_4, 64_ = 0.221, p = 0.926; Figure S3).

## Supplementary Materials

**Figure S1.**
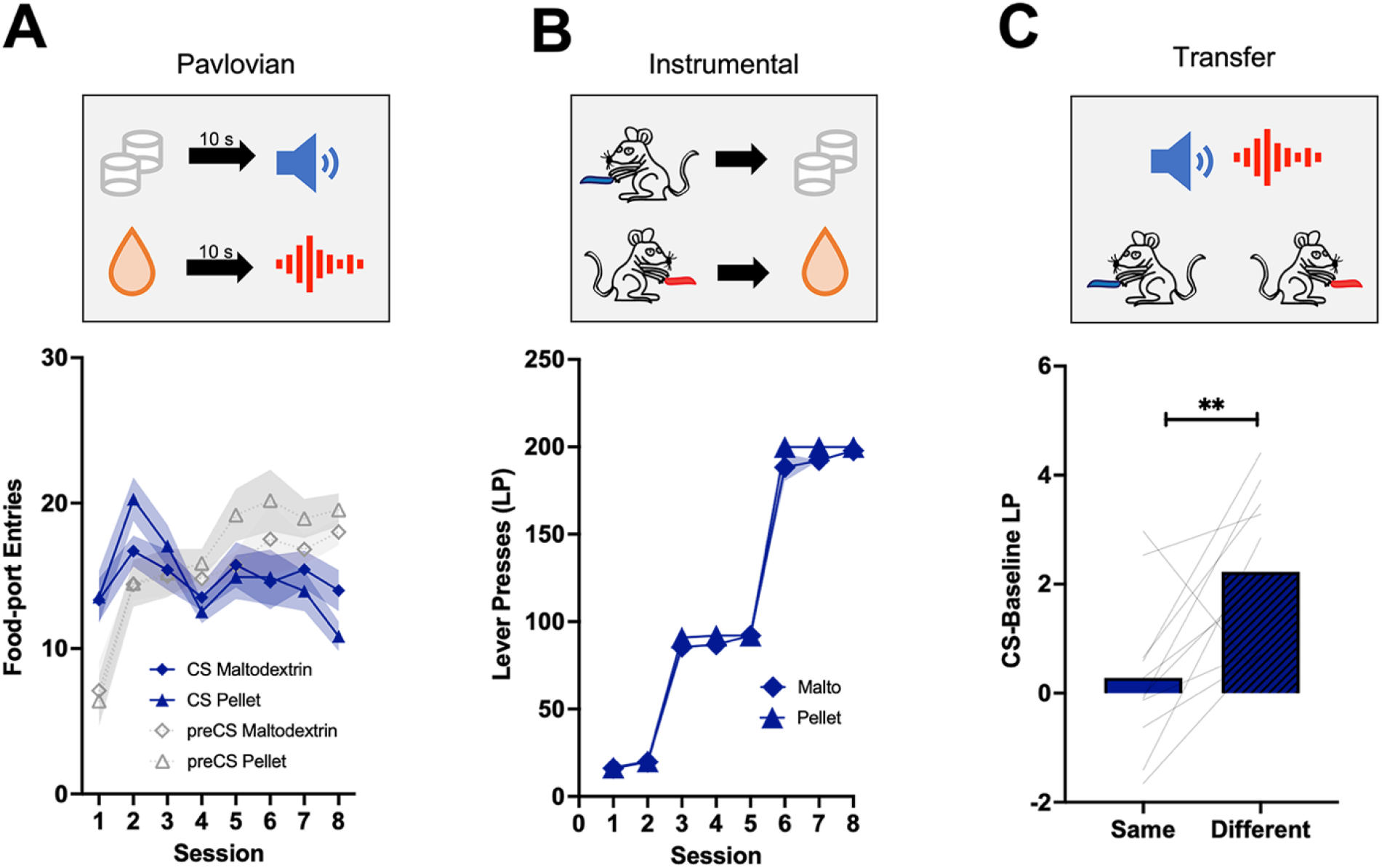
Backward conditioning Pavlovian to Instrumental Transfer Test Pilot study. **A)** Backward Pavlovian training. **B)** Instrumental conditioning. **C)** Transfer test.

**Figure S2.**
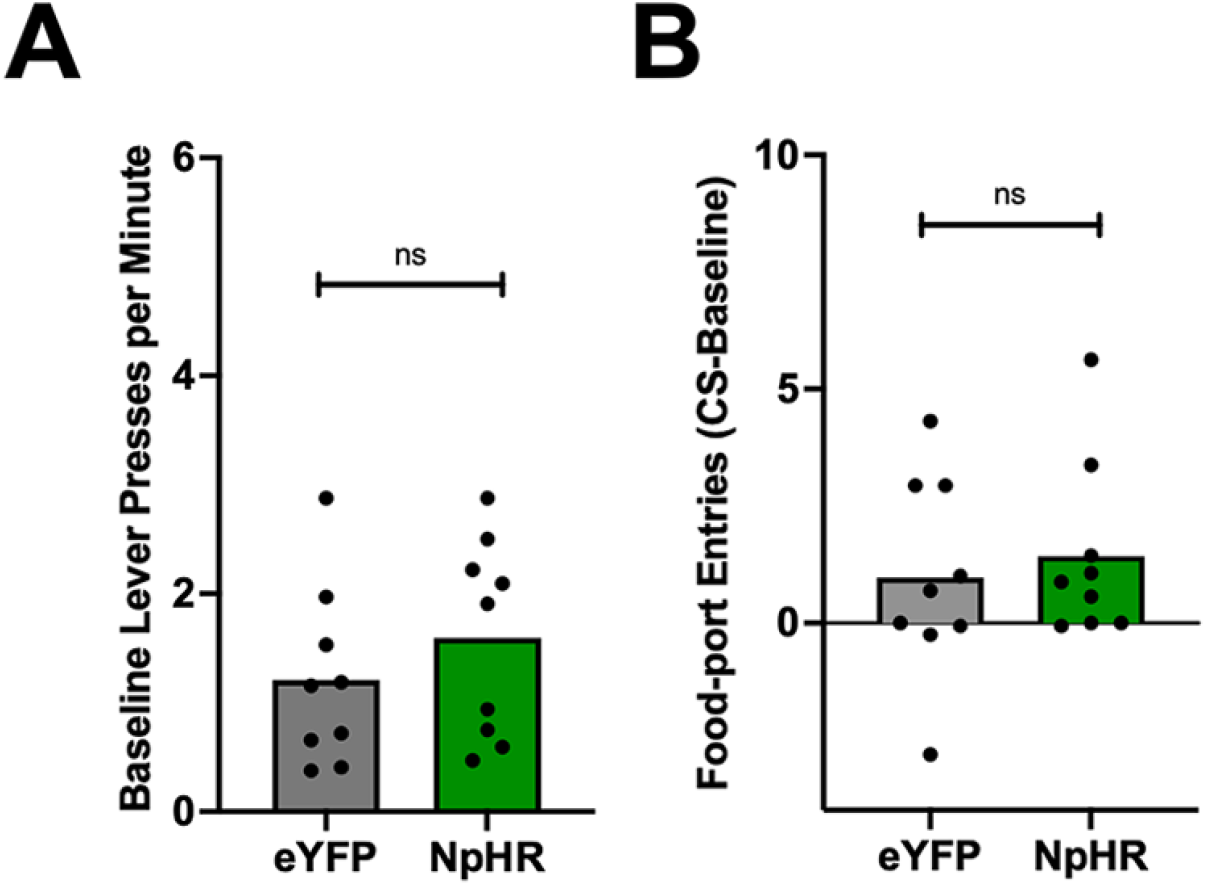
No difference in baseline lever pressing or head entry responses during transfer test. **A)** Transfer test data are displayed in Figure 2C as lever presses made relative to baseline responding. Those baseline levels are shown here and do not differ between groups, t_16_ = 0.946, p = 0.358. **B)** During the transfer test rats also had the ability to enter the food port as well as lever press. We find the backward cues have little effect on head entries into the food port in both groups, t_16_ = 0.946, p = 0.358.

**Figure S3.**
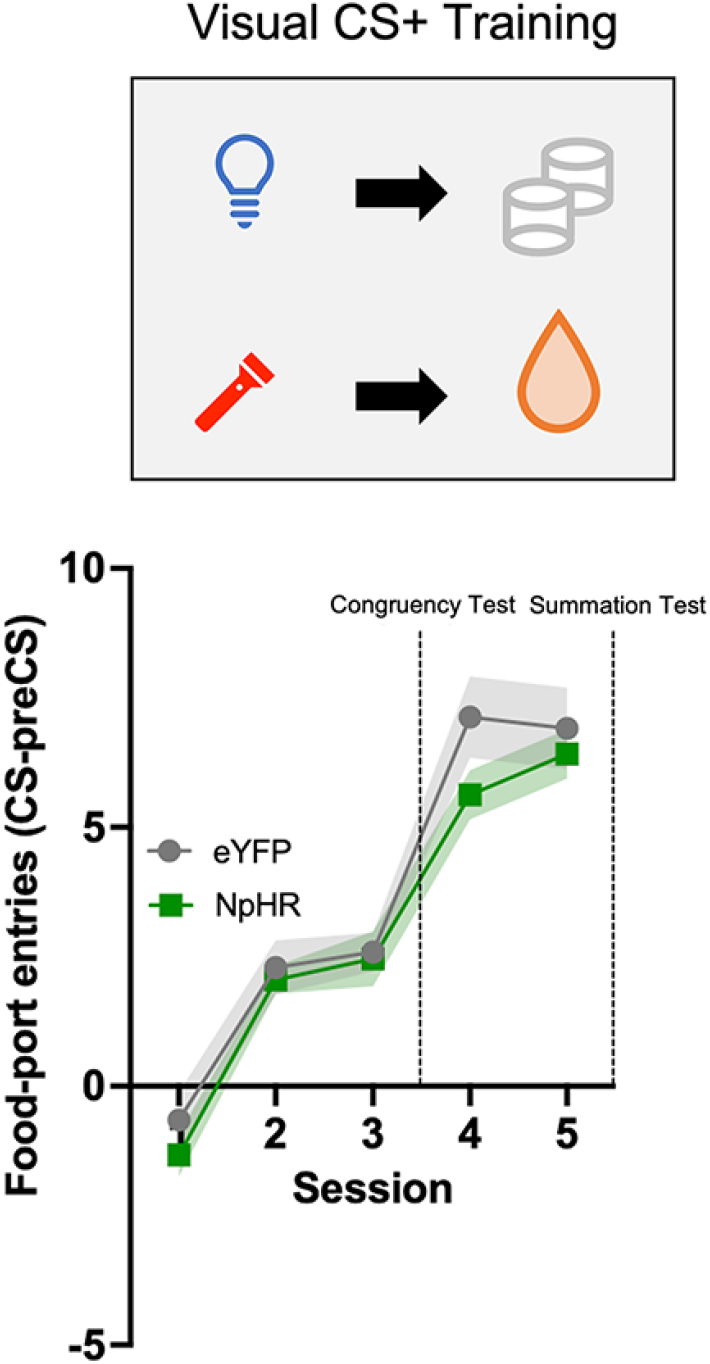
Forward conditioning with visual cues. Rats learned relationships between two visual cues (house light and flashing-light) and two rewards (sucrose pellet and maltodextrin solution). Rats received 3 days of this training before completing the Congruency test and then two more days of training before the Summation test.

## Notes

### Competing Interest Statement

The authors have declared no competing interest.

## References

1. Pavlov, I.P. (1927). Conditioned reflexes: An investigation of the physiological activity of the cerebral cortex (Oxford University Press).

2. Bolles, R.C. (1993). The story of psychology: a thematic history (Brooks/Cole Publishing Co.).

3. Rescorla, R.A., and Wagner, A.R. (1972). A Theory of Pavlovian Conditioning: Variations in the Effectiveness of Reinforcement and Nonreinforcement. In Classical conditioning II: current research and theory, A. Black and W. Prokasy, eds. (Appleton-Centrury-Crofts), pp. 64–99.

4. Mackintosh, N.J. (1975). A theory of attention: Variations in the associability of stimuli with reinforcement. Psychol. Rev. 82, 276–298.

5. Sutton, R.S., and Barto, A.G. (1981). Toward a modern theory of adaptive networks: Expectation and prediction. Psychol. Rev. 88, 135–170.

6. Pearce, J.M., and Hall, G. (1980). A model for Pavlovian learning: Variations in the effectiveness of conditioned but not of unconditioned stimuli. Psychol. Rev. 87, 532–552.

7. Kamin, L.J. (1969). Predictability, Surprise, Attention, and Conditioning. In Punishment Aversive Behavior, B. A. Campbell and R.M. Church, eds. (Appleton-Century Crofts.), pp. 279–296.

8. Daw, N.D., Niv, Y., and Dayan, P. (2005). Uncertainty-based competition between prefrontal and dorsolateral striatal systems for behavioral control. Nat. Neurosci. 2005 812 8, 1704–1711.

9. Clark, A. (2013). Whatever next? Predictive brains, situated agents, and the future of cognitive science. Behav. Brain Sci. 36, 181–204.

10. Schultz, W., and Dickinson, A. (2003). Neuronal Coding of Prediction Errors. Annu. Rev. Neurosci. 23, 473–500.

11. Miller, R.R., Barnet, R.C., and Grahame, N.J. (1995). Assessment of the Rescorla-Wagner model. Psychol. Bull. 117, 363–386.

12. Schultz, W., Dayan, P., and Montague, P.R. (1997). A neural substrate of prediction and reward. Science (80-.). 275, 1593–1599.

13. Glimcher, P.W. (2011). Understanding dopamine and reinforcement learning: The dopamine reward prediction error hypothesis. Proc. Natl. Acad. Sci. U. S. A. 108, 15647–15654.

14. Waelti, P., Dickinson, A., and Schultz, W. (2001). Dopamine responses comply with basic assumptions of formal learning theory. Nature 412, 43–48.

15. Schultz, W. (2016). Dopamine reward prediction-error signalling: A two-component response. Nat. Rev. Neurosci. 17, 183–195.

16. Cohen, J.Y., Haesler, S., Vong, L., Lowell, B.B., and Uchida, N. (2012). Neuron-type-specific signals for reward and punishment in the ventral tegmental area. Nature 482, 85–88.

17. Saunders, B.T., Richard, J.M., Margolis, E.B., and Janak, P.H. (2018). Dopamine neurons create Pavlovian conditioned stimuli with circuit-defined motivational properties. Nat. Neurosci. 2018 218 21, 1072–1083.

18. Hollerman, J.R., and Schultz, W. (1998). Dopamine neurons report an error in the temporal prediction of reward during learning. Nat. Neurosci. 1, 304–309.

19. Chang, C.Y., Esber, G.R., Marrero-Garcia, Y., Yau, H.J., Bonci, A., and Schoenbaum, G. (2015). Brief optogenetic inhibition of dopamine neurons mimics endogenous negative reward prediction errors. Nat. Neurosci. 2016 191 19, 111–116.

20. Steinberg, E.E., Keiflin, R., Boivin, J.R., Witten, I.B., Deisseroth, K., and Janak, P.H. (2013). A causal link between prediction errors, dopamine neurons and learning. Nat. Neurosci. 16, 966–973.

21. Tsai, H.C., Zhang, F., Adamantidis, A., Stuber, G.D., Bond, A., De Lecea, L., and Deisseroth, K. (2009). Phasic firing in dopaminergic neurons is sufficient for behavioral conditioning. Science 324, 1080–1084.

22. Lak, A., Stauffer, W.R., and Schultz, W. (2014). Dopamine prediction error responses integrate subjective value from different reward dimensions. Proc. Natl. Acad. Sci. U. S. A. 111, 2343–2348.

23. Tobler, P.N., Fiorillo, C.D., and Schultz, W. (2005). Adaptive coding of reward value by dopamine neurons. Science 307, 1642–1645.

24. Fiorillo, C.D. (2013). Two dimensions of value: dopamine neurons represent reward but not aversiveness. Science 341, 546–549.

25. Stalnaker, T.A., Howard, J.D., Takahashi, Y.K., Gershman, S.J., Kahnt, T., and Schoenbaum, G. (2019). Dopamine neuron ensembles signal the content of sensory prediction errors. Elife 8.

26. Howard, J.D., and Kahnt, T. (2018). Identity prediction errors in the human midbrain update reward-identity expectations in the orbitofrontal cortex. Nat. Commun. 2018 91 9, 1–11.

27. Sharpe, M.J., Batchelor, H.M., Mueller, L.E., Yun Chang, C., Maes, E.J.P., Niv, Y., and Schoenbaum, G. (2020). Dopamine transients do not act as model-free prediction errors during associative learning. Nat. Commun. 11, 1–10.

28. Sadacca, B.F., Jones, J.L., and Schoenbaum, G. (2016). Midbrain dopamine neurons compute inferred and cached value prediction errors in a common framework. Elife 5.

29. Takahashi, Y.K., Batchelor, H.M., Liu, B., Khanna, A., Morales, M., and Schoenbaum, G. (2017). Dopamine Neurons Respond to Errors in the Prediction of Sensory Features of Expected Rewards. Neuron 95, 1395–1405.e3.

30. Chang, C.Y., Gardner, M., Di Tillio, M.G., and Schoenbaum, G. (2017). Optogenetic Blockade of Dopamine Transients Prevents Learning Induced by Changes in Reward Features. Curr. Biol. 27, 3480–3486.e3.

31. Sharpe, M.J., Chang, C.Y., Liu, M.A., Batchelor, H.M., Mueller, L.E., Jones, J.L., Niv, Y., and Schoenbaum, G. (2017). Dopamine transients are sufficient and necessary for acquisition of model-based associations. Nat. Neurosci. 20, 735–742.

32. Keiflin, R., Pribut, H.J., Shah, N.B., and Janak, P.H. (2019). Ventral Tegmental Dopamine Neurons Participate in Reward Identity Predictions. Curr. Biol. 29, 93–103.e3.

33. Engelhard, B., Finkelstein, J., Cox, J., Fleming, W., Jang, H.J., Ornelas, S., Koay, S.A., Thiberge, S.Y., Daw, N.D., Tank, D.W., et al. (2019). Specialized coding of sensory, motor and cognitive variables in VTA dopamine neurons. Nat. 2019 5707762 570, 509–513.

34. Barnet, R.C., and Miller, R.R. (1996). Scond-order excitation mediated by a backward conditioned inhibitor. J. Exp. Psychol. Anim. Behav. Process. 22, 279–296.

35. Prével, A., Rivière, V., Darcheville, J.C., Urcelay, G.P., and Miller, R.R. (2019). Excitatory second-order conditioning using a backward first-order conditioned stimulus: A challenge for prediction error reduction. Q. J. Exp. Psychol. 72, 1453–1465.

36. Chang, R.C., Blaisdell, A.P., and Miller, R.R. (2003). Backward Conditioning: Mediation by the Context. J. Exp. Psychol. Anim. Behav. Process. 29, 171–183.

37. Urushihara, K. (2004). Excitatory backward conditioning in an appetitive conditioned reinforcement preparation with rats. Behav. Processes 67, 477–489.

38. Cole, R.P., and Miller, R.R. (1999). Conditioned Excitation and Conditioned Inhibition Acquired through Backward Conditioning. Learn. Motiv. 30, 129–156.

39. Laurent, V., Wong, F.L., and Balleine, B.W. (2017). The lateral habenula and its input to the rostromedial tegmental nucleus mediates outcome-specific conditioned inhibition. J. Neurosci. 37, 10932–10942.

40. Laurent, V., and Balleine, B.W. (2015). Factual and counterfactual action-outcome mappings control choice between goal-directed actions in rats. Curr. Biol. 25, 1074–1079.

41. Laurent, V., Wong, F.L., and Balleine, B.W. (2015). δ-Opioid receptors in the accumbens shell mediate the influence of both excitatory and inhibitory predictions on choice. Br. J. Pharmacol. 172, 562–570.

42. Witten, I.B., Steinberg, E.E., Lee, S.Y., Davidson, T.J., Zalocusky, K.A., Brodsky, M., Yizhar, O., Cho, S.L., Gong, S., Ramakrishnan, C., et al. (2011). Recombinase-driver rat lines: Tools, techniques, and optogenetic application to dopamine-mediated reinforcement. Neuron 72, 721–733.

43. Maes, E.J.P., Sharpe, M.J., Usypchuk, A.A., Lozzi, M., Chang, C.Y., Gardner, M.P.H., Schoenbaum, G., and Iordanova, M.D. (2020). Causal evidence supporting the proposal that dopamine transients function as temporal difference prediction errors. Nat. Neurosci. 23, 176–178.

44. Morrens, J., Aydin, Ç., Janse van Rensburg, A., Esquivelzeta Rabell, J., and Haesler, S. (2020). Cue-Evoked Dopamine Promotes Conditioned Responding during Learning. Neuron 106, 142–153.e7.

45. Schultz, W. (1998). Predictive reward signal of dopamine neurons. J. Neurophysiol. 80, 1–27.

46. Langdon, A.J., Sharpe, M.J., Schoenbaum, G., and Niv, Y. (2018). Model-based predictions for dopamine. Curr. Opin. Neurobiol. 49, 1–7.

47. Sharpe, M.J., and Schoenbaum, G. (2018). Evaluation of the hypothesis that phasic dopamine constitutes a cached-value signal. Neurobiol. Learn. Mem. 153, 131–136.

48. Sharpe, M.J., Batchelor, H.M., and Schoenbaum, G. (2017). Preconditioned cues have no value. Elife 6.

49. Keiflin, R., Pribut, H.J., Shah, N.B., and Janak, P.H. (2019). Ventral Tegmental Dopamine Neurons Participate in Reward Identity Predictions. Curr. Biol. 29, 93–103.e3.

50. Gardner, M.P.H., Schoenbaum, G., and Gershman, S.J. (2018). Rethinking dopamine as generalized prediction error. Proc. R. Soc. B 285.

51. Seitz, B.M., Blaisdell, A.P., and Sharpe, M.J. (2021). Higher-Order Conditioning and Dopamine: Charting a Path Forward. Front. Behav. Neurosci. 0, 228.

52. Millard, S.J., Bearden, C.E., Karlsgodt, K.H., and Sharpe, M.J. (2021). The prediction-error hypothesis of schizophrenia: new data point to circuit-specific changes in dopamine activity. Neuropsychopharmacol. 2021, 1–13.

53. Jensen, J., Willeit, M., Zipursky, R.B., Savina, I., Smith, A.J., Menon, M., Crawley, A.P., and Kapur, S. (2007). The Formation of Abnormal Associations in Schizophrenia: Neural and Behavioral Evidence. Neuropsychopharmacol. 2008 333 33, 473–479.

54. Corlett, P.R., Taylor, J.R., Wang, X.J., Fletcher, P.C., and Krystal, J.H. (2010). Toward a Neurobiology of Delusions. Prog. Neurobiol. 92, 345.

55. Corlett, P.R., Murray, G.K., Honey, G.D., Aitken, M.R.F., Shanks, D.R., Robbins, T.W., Bullmore, E.T., Dickinson, A., and Fletcher, P.C. (2007). Disrupted prediction-error signal in psychosis: evidence for an associative account of delusions. Brain 130, 2387–2400.

56. Morris, R.W., Vercammen, A., Lenroot, R., Moore, L., Langton, J.M., Short, B., Kulkarni, J., Curtis, J., O’Donnell, M., Weickert, C.S., et al. (2012). Disambiguating ventral striatum fMRI-related BOLD signal during reward prediction in schizophrenia. Mol. Psychiatry 17, 280–289.

57. Morris, R., Griffiths, O., Le Pelley, M.E., and Weickert, T.W. (2013). Attention to irrelevant cues is related to positive symptoms in schizophrenia. Schizophr. Bull. 39, 575–582.

58. Hoang, I.B., and Sharpe, M.J. (2021). The basolateral amygdala and lateral hypothalamus bias learning towards motivationally significant events. Curr. Opin. Behav. Sci. 41, 92–97.

59. Sharpe, M.J., Batchelor, H.M., Mueller, L.E., Gardner, M.P.H., and Schoenbaum, G. (2021). Past experience shapes the neural circuits recruited for future learning. Nat. Neurosci. 2021 243 24, 391–400.

